# Gene duplicates cause hybrid lethality between sympatric species of *Mimulus*

**DOI:** 10.1101/201392

**Authors:** Matthew P. Zuellig, Andrea L. Sweigart

## Abstract

Hybrid incompatibilities play a critical role in the evolution and maintenance of species. We have discovered a simple genetic incompatibility that causes lethality in hybrids between two closely related species of yellow monkeyflower (*Mimulus guttatus* and *M. nasutus*). This hybrid incompatibility, which causes one sixteenth of F_2_ hybrid seedlings to lack chlorophyll and die shortly after germination, occurs between sympatric populations that are connected by ongoing interspecific gene flow. Using complimentary genetic mapping and gene expression analyses, we show that lethality occurs in hybrids that lack a functional copy of the critical photosynthetic gene *pTAC14*. In *M. guttatus*, this gene was duplicated, but the ancestral copy is no longer expressed. In *M. nasutus*, the duplication is missing altogether. As a result, hybrids die when they are homozygous for the nonfunctional *M. guttatus* copy and missing the duplicate from *M. nasutus*, apparently due to misregulated transcription of key photosynthetic genes. Our study indicates that neutral evolutionary processes may play an important role in the evolution of hybrid incompatibilities and opens the door to direct investigations of their contribution to reproductive isolation among naturally hybridizing species.

**Author Summary:** Hybrid incompatibilities play an important role in speciation, because they act to limit gene flow between species. Identifying the genes that underlie these barriers sheds light on the evolutionary forces and genetic mechanisms that give rise to new species. We identified a reproductive barrier that causes lethality in the F2 offspring of sympatric species of yellow monkeyflower (*Mimulus guttatus* and *M. nasutus*). We show that lethality occurs in hybrids that lack a functional copy of the critical photosynthetic gene *pTAC14*. This gene was duplicated in *M. guttatus*, but the ancestral copy subsequently lost function. In *M. nasutus*, no duplication occurred. As a consequence, F2 hybrids that are homozygous for non-functional *M. guttatus* copies at one locus and missing *M. nasutus* duplicates at the other locus completely lack functional *pTAC14* and die. Our data indicate that non-functionalization of ancestral *pTAC14* in *M. guttatus* occurred via neutral evolutionary change. These results suggest that neutral evolutionary forces may play an important role in speciation.

## Introduction

Across diverse taxa, hybrid incompatibilities arise as a byproduct of genetic divergence among incipient species. The basic genetic underpinnings of this process are well understood: two or more mutational differences between species interact epistatically to cause hybrid inviability or sterility [1-3]. However, what is less clear, and often very challenging to uncover, is the nature of the molecular changes and evolutionary forces that lead to hybrid incompatibilities. What sort of mutations are perfectly functional within species but cause reproductive failure or death in hybrids? Do such mutations accumulate within species by neutral processes or are they positively selected, perhaps providing an ecological advantage or resolving an intragenomic conflict? Addressing the first of these questions is most straightforward in systems with well-developed genetic tools that facilitate positional cloning, which explains why most progress has been made in traditional models like *Drosophila*, *Arabidopsis*, and rice. However, insight into the evolutionary forces acting on hybrid incompatibilities during the speciation process requires a focus on young species pairs with natural populations.

Over the past two decades, genetic dissection of diverse incompatibilities has provided some hints about their evolutionary origins (reviewed in [4-7]). Often, hybrid incompatibility genes show molecular signatures of positive selection [8-13], and there is suggestive evidence that incompatibility alleles can arise through ecological adaptation [14-16] or recurrent bouts of intragenomic conflict [11, 17-19]. On the other hand, there is also evidence from a handful of cases, all involving gene duplicates, that the evolution of hybrid dysfunction need not involve natural selection [20-22].

The idea that gene duplication might play a key role in hybrid incompatibilities was initially proposed by Muller as a variant of his original model (1942). He explained how gene duplication, followed by degenerative mutations and divergent copy loss, could lead to a difference in gene position between species with missing (or inactive) copies acting as recessive incompatibility alleles [3]. This same scenario was emphasized later as an explanation for defects in pollen development between subspecies of Asian cultivated rice, *Oryza sativa* [23], and more recently, as a general mechanism of hybrid breakdown via neutral processes [24, 25]. There have now been three empirical demonstrations of gene transposition giving rise to interspecific hybrid male sterility; one case involves *Drosophila melanogaster*-*D. simulans* hybrids [26] and the other two arise from crosses between *O. sativa* and wild species [21, 22]. Gene duplication/transposition also causes lethal and sterile combinations that segregate within *Arabidopsis thaliana* [27, 28] and *O. sativa* [20]. However, it is not yet clear whether divergent resolution of gene duplicates contributes to hybrid incompatibilities between wild species in the early stages of divergence. Only by identifying examples in young species pairs, particularly those with sympatric populations and still connected by some degree of gene flow, will it be possible to evaluate the contribution of such loci to speciation.

In this study we investigate the molecular genetic basis of hybrid seedling lethality between two closely related species of yellow monkeyflower, *Mimulus guttatus* and *M. nasutus*. These recently diverged species (200-500kya; [29]) co-occur throughout much of their shared range in western North America, where reproductive isolation between sympatric populations occurs through a number of prezygotic [30-33] and postzygotic barriers [34-40]. Despite substantial reproductive isolation, patterns of shared variation across their genomes indicate historical and ongoing gene flow between the two species [29, 31, 41]. Here we focus on sympatric populations of *M. guttatus* and *M. nasutus* located at Don Pedro Reservoir (DPR) in central California, where both species coexist within centimeters of one another. Species at DPR are strongly isolated by divergence in flowering time and mating system [33]; nevertheless, studies have shown low levels of hybridization [33] and a clear signal of introgression [29]. Using high-resolution genetic mapping and genome-wide expression analyses we identify a duplicate gene pair as the cause of *Mimulus* hybrid lethality. As the first case of hybrid incompatibility genes identified between naturally hybridizing species, this study opens the door to direct investigations of their evolutionary dynamics and contribution to reproductive isolation.

## Results

### Hybrid lethality is caused by a two-locus incompatibility

Hybrid lethality occurs in the hybrid progeny DPRG102 and DPRN104 and is easily characterized by seedlings that completely lack chlorophyll (white seedlings, see Fig S1). As a first step toward investigating the genetic basis of hybrid lethality between inbred lines of *M. guttatus* (DPRG102) and *M. nasutus* (DPRN104) from the sympatric DPR site, we examined phenotypic ratios of white and green seedlings among their selfed progeny and reciprocal F1 and F2 hybrids (Fig S1, Table S1). Although we never observed white seedlings in the selfed progeny of parental lines or in F1 hybrids, we discovered that roughly 1/16 of F2 hybrid seedlings were white (maternal parent listed first: DPRG102xDPRN104, *N*=516, 7.36% white seedlings; DPRN104xDPRG102, *N*=661, 5.75% white seedlings). Segregation of white seedlings in reciprocal F2 hybrids suggests a nuclear, rather than cyto-nuclear, genetic incompatibility. Chi-squared tests rejected several genetic models that could potentially explain the observed phenotypic ratios, but could not reject a two-locus model involving only recessive alleles in either F2 population, or when their ratios were combined (Table S1). These results suggest that hybrid lethality between sympatric *M. guttatus* and *M. nasutus* is caused by a two-locus, recessive-recessive hybrid incompatibility.

To genetically map *Mimulus* hybrid lethality, we performed two rounds of bulked segregant analysis (BSA). In the first round, we pooled DNA from green and white F2 seedlings into eight separate tubes (six individuals per pool, four replicates each for green and white). Because incompatibility alleles act recessively, our expectation was that pooled white seedlings should be homozygous (for either DPRG102 or DPRN104 alleles) at markers linked to hybrid lethality loci, whereas green seedlings should segregate 1:2:1 (for DPRG102 homozygotes: heterozygotes: DPRN104 homozygotes). Of the 126 size-polymorphic markers (spanning much of the *Mimulus* genome) that we used for genotyping, four showed an association with seedling phenotype: the four tubes with white seedlings carried only (or mostly) DPRN104 alleles, whereas green seedlings carried both parental alleles. All four markers map to a region of roughly 40 cM on linkage group 14 (inferred by marker position in Fishman et al. 2014), which we named *hybrid lethal 14* (*hl14*).

To identify the partner locus, we performed a second round of BSA controlling for genotype at *hl14*. We generated 60 F3 families by self-fertilizing green F2 hybrids that were homozygous for DPRN104 alleles at *hl14* (determined by genotyping flanking markers); these F3 families segregated green and white seedlings in ratios of either 3:1 or 1:0. We reasoned that if hybrid lethality is caused by *hl14* and a single interacting locus, white F3 hybrids should be homozygous for DPRG102 alleles at the partner, whereas green F3 families that do not segregate white seedlings should be homozygous for DPRN104 alleles. Based on this logic, we formed two separate pools of DNA from F3 hybrids: one with 34 white seedlings and one with 26 green seedlings from non-segregating families. Note that each F3 seedling was derived from a different family (i.e., from a unique F2 maternal parent) so that at markers unlinked to hybrid lethality, both pools should carry each of the two parental alleles at ~50% frequency. For the two pools, we performed whole genome sequencing, generated a genome-wide SNP dataset, and calculated average allele frequency difference in 200-SNP sliding windows (100-SNP overlap between windows). Using this approach, we discovered that the top 5% most divergent windows were located in contiguous windows along the distal end of chromosome 13 (Fig S2), which we named *hybrid lethal 13* (*hl13*).

To fine-map *hl13* and *hl14*, we generated a large DPRN104 × DPRG102 F2 mapping population, oversampling white seedlings to roughly equalize frequencies of the two phenotypes (white = 44%, green = 56%, *N* = 2,652). Each F2 individual was genotyped at markers believed to flank the hybrid lethality loci: M208 and M236 at *hl13*, and M241 and M132 *hl14*. As expected, nearly all white seedlings were homozygous for DPRG102 alleles at the hl13-linked markers and homozygous for DPRN104 alleles at the *hl14* markers (92%, *N* = 1174), whereas green seedlings never carried this genotype (*N* = 1478) (Fig 1). Because we later discovered that both M208 and M236 are proximal to *hl13*, an additional 2,182 F2 hybrids were screened with a more distal marker (either M263 or M255). We genotyped informative recombinants at additional size-polymorphic and SNP-based markers designed in each interval. Although white seedlings must be destructively sampled for DNA, green seedlings were allowed to grow into adult plants so that informative recombinants could be self-fertilized and phenotyped via progeny testing. In this way, we determined if green F2 hybrids were heterozygous for *hl13* and/or *hl14* (versus homozygous for compatible alleles). Using this strategy, we mapped the *hl13* locus to a 72.2 kb-region at the distal end of chromosome 13 that contains 24 genes (Fig 2A). At the same time, we mapped the *hl14* locus to a 51.6 kb-region of chromosome 14 that contains six genes, as well as a gap of unknown size in the *M. guttatus* IM62 reference genome (Fig 2B). For each *hl13* and *hl14* candidate gene, we identified its top blast hit(s) in *Arabidopsis thaliana*, gene ontology terms, known mutant phenotypes, and predicted functions (Table S2).

**Fig 1:**
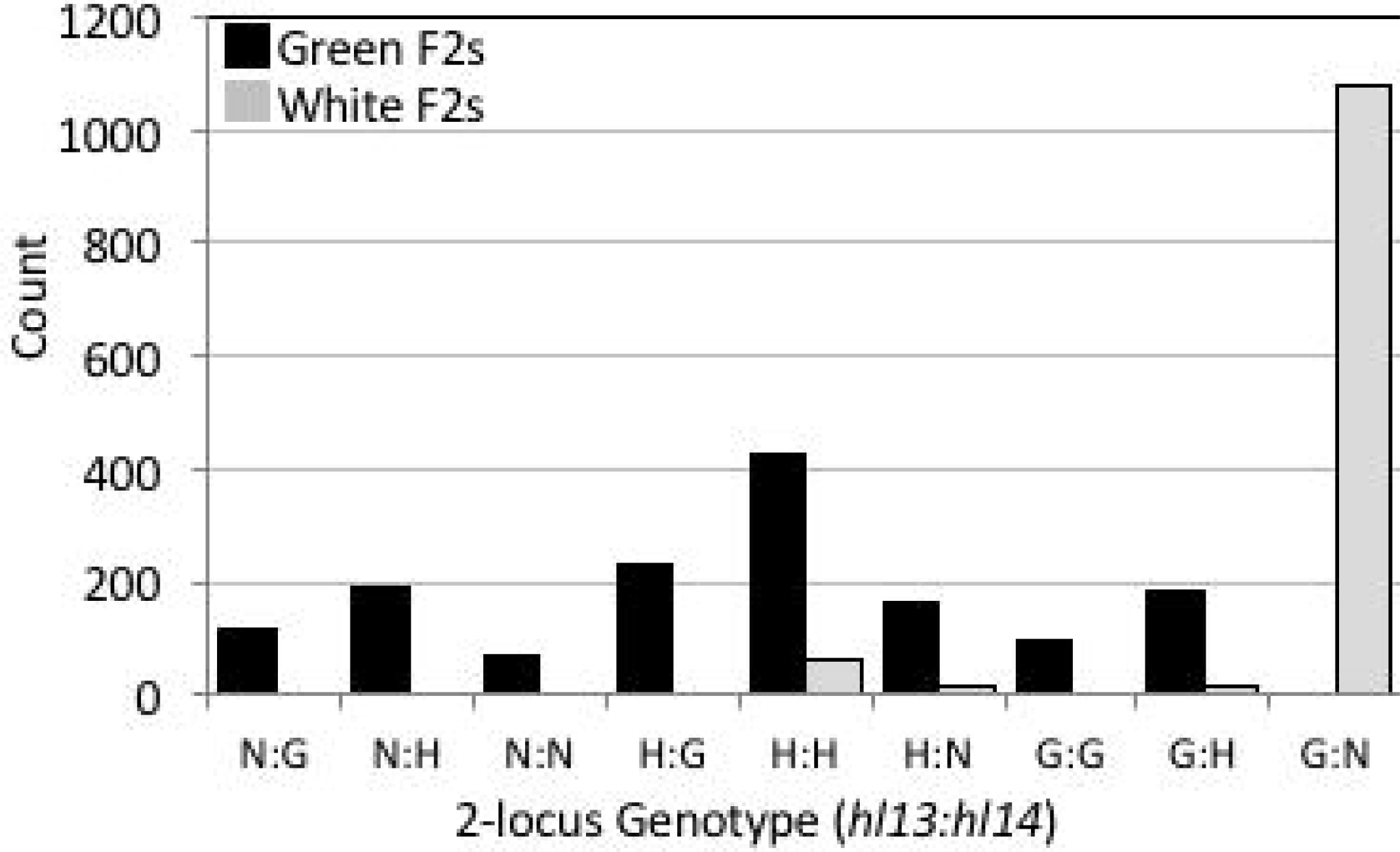
Two-locus genotypes of green and white seedlings at markers linked to *hl13* and *hl14*. Count represents the number of F2s with a given genotype. Genotypes are homozygous *M. guttatus* (‘G’), homozygous *M. nasutus* (‘N’), and heterozygous (‘H’). The vast majority (92%) of white F2s carry the G:N genotype, whereas green F2s carry all genotypes except G:N. Sample size is 2,652, which represents the subset of our mapping population that was genotyped with markers M208, M236, M241, and M132.

**Fig 2:**
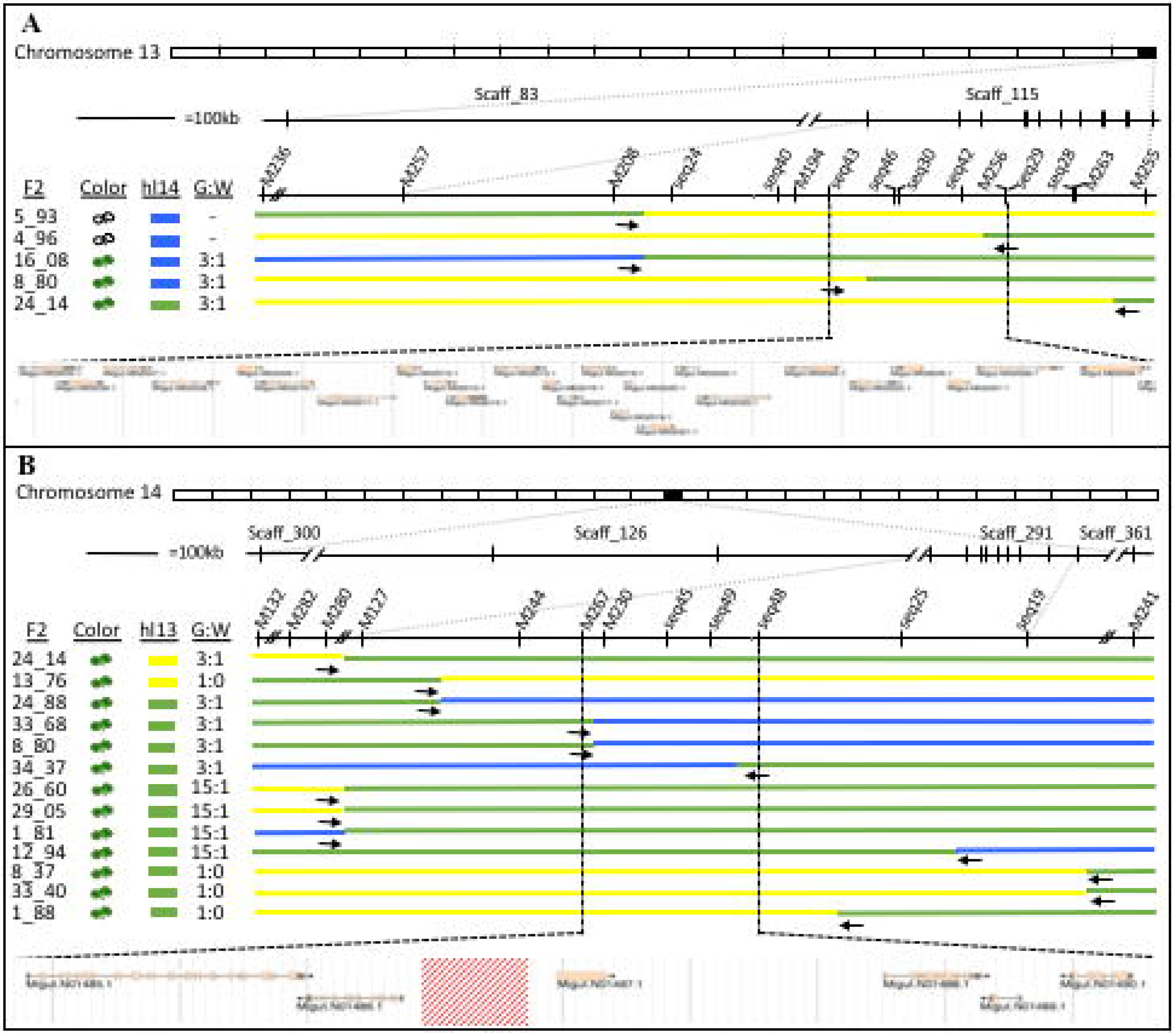
Fine-mapping localizes hybrid lethality loci to small genomic intervals. (A) *hl13* maps to a 72kb region on scaffold 115 containing 22 genes. (B) *hl14* maps to a 51.6kb region on scaffold 291 containing 6 genes and a gap in the reference genome (red box) of unknown size. Horizontal bars represent F2 recombinants that were informative for mapping *hl13* (between M208 and M255) and *hl14* (between M280 and M241), where marker genotypes are yellow (homozygous for DPRG102 alleles), blue (homozygous for DPRN104 alleles), and green (heterozygous). F2 individual used for mapping, F2 phenotype (white or green), genotype at partner locus (DPRG102, DPRN104, or heterozygous), and green:white ratio of F3 progeny are given for each recombinant. Vertical and diagonal hatch marks are marker positions and breaks within/between scaffolds, respectively. Scaffolds from v1.1 of the reference genome are included in figure, though all gene annotation and naming is based on the updated v2.0 assembly (phytozome.org).

### Gene duplicates map to hybrid lethality loci

Among several strong functional candidates for *hl13* is *Migut.M02023*, a homolog of *pTAC14* (*PLASTID TRANSCRIPTIONALLY ACTIVE CHROMOSOME 14*). In *A. thaliana*, *pTAC14* is essential for proper chloroplast development and mutants show a chlorotic lethal phenotype [42] that appears identical to DPRG102**×**DPRN104 F2 hybrid lethality. In addition to *Migut.M02023* on chromosome 13, we also identified a highly similar and slightly truncated protein homolog of *pTAC14* (99.1% amino acid similarity along length of truncated homolog), *Migut.O00467*, located on an unmapped scaffold of the IM62 *M. guttatus* reference genome (v2.0 scaffold_193). To investigate the possibility that this additional copy of *Mg.pTAC14* resides on chromosome 14, we turned to several large-insert IM62 genomic libraries (six fosmid and two BAC libraries) that were generated and end-sequenced as part of the reference genome assembly effort [43]. Among these libraries, only a single end-sequence of one fosmid blasts to v2.0 scaffold_193. Intriguingly, the other end-sequence of this same fosmid blasts to the first exon of *Migut.N01489*, a gene within the mapped interval of *hl14*. This finding provides evidence that a second copy of *Mg.pTAC14* is located on chromosome 14 in IM62, despite it being absent from the current genome assembly. Using PCR, we confirmed that the DPRG102 genome also contains two copies of *pTAC14* (Fig S3). However, despite exhaustive PCR and cloning efforts (using many different primer combinations), we recovered only one copy of *pTAC14* from DPRN104 genomic DNA.

To determine if *Mimulus pTAC14* duplicates genetically map to DPRG102-DPRN104 hybrid lethality loci, we obtained a set of 96 DPRG102xDPRN104 F2 hybrids carrying each of the nine possible two-locus genotypes at *hl13* and *hl14* (10 replicates for each green genotype and 16 replicates of the white seedling genotype, see Fig 3). Using a set of conserved primers spanning exons 6-8, we PCR-amplified and sequenced *Mimulus pTAC14* from each of these F2 hybrids. Across this region, 10 SNPs define three distinct haplotypes of *pTAC14*: “G1” and “G2” from DPRG102 and “N1” DPRN104 (Fig 3). Remarkably, we discovered a perfect association between *pTAC14* haplotype and *hl13/hl14* genotype: G1 is present in all individuals with DPRG102 alleles at *hl13*, G2 is in all individuals with DPRG102 alleles at *hl14*, and N1 is in all individuals with DPRN104 alleles at *hl13*. From this pattern, we infer that both DPRG102 and DPRN104 carry copies of *pTAC14* at *hl13* (hereafter referred to as *Mg.pTAC14*_*1* and *Mn.pTAC14*_*1*, respectively), but that only DPRG102 carries a copy at *hl14* (referred to as *Mg.pTAC14*_*2*).

**Fig 3:**
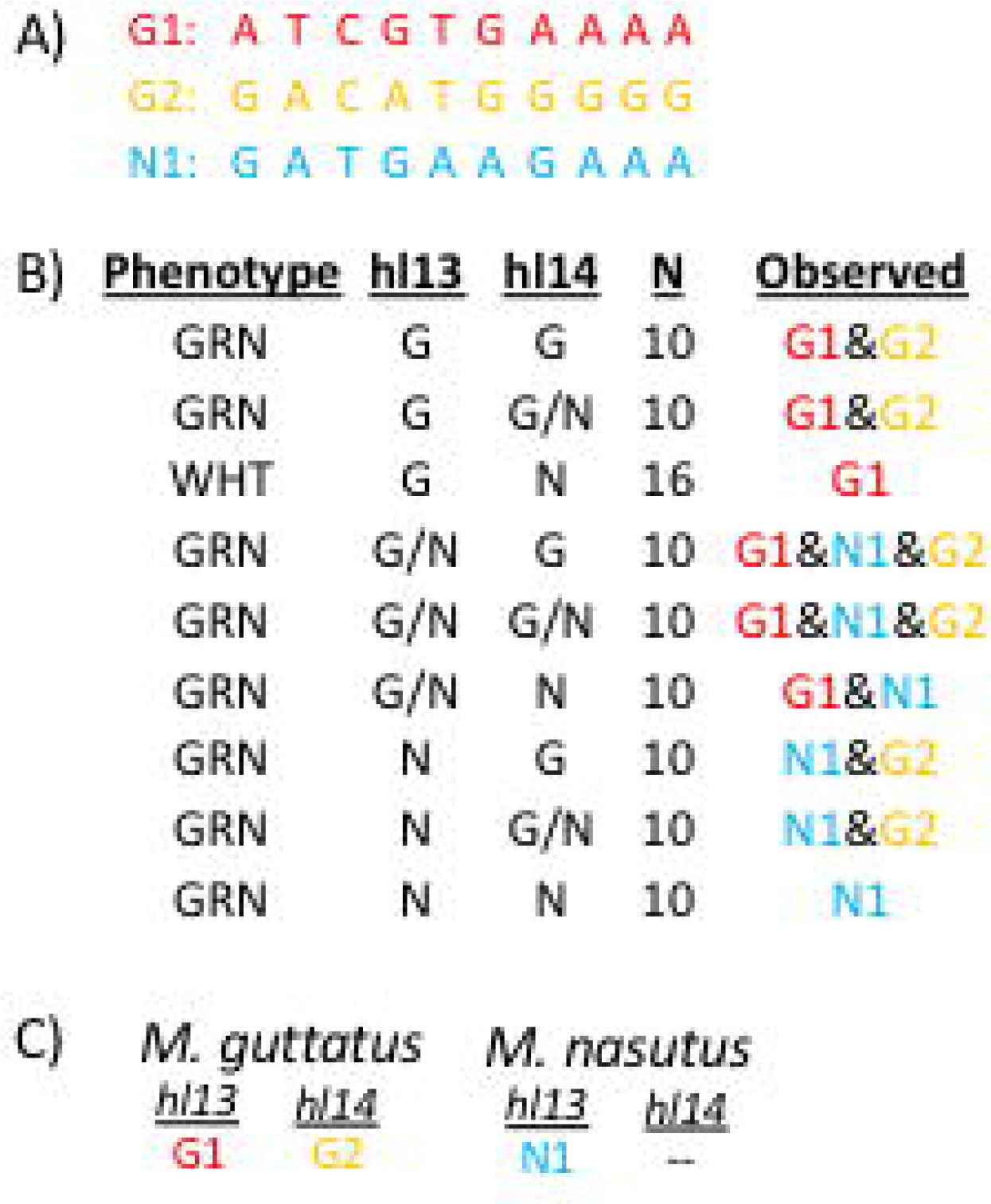
Duplicate copies of pTAC14 map to both hybrid lethality loci in *M. guttatus*. (A) Ten SNPs differentiate Mg.pTAC14_1 (G1), Mg.pTAC14_2 (G2), and Mnas.pTAC14 (N1). (B) Phenotype: green (GRN) or white (WHT) F2. hl13 and hl14: genotype at flanking markers [G (DPRG102), N (DPRN104, and G/N (heterozygous)]. N is sample size. Observed: Copies of *pTAC14* observed in each F2, consistent across all F2s for a given genotype. Note that individuals with DPRG102 alleles at hl13 always carry Mg.pTAC14_1, individuals with DPRG102 alleles at hl14 always carry Mg.pTAC14_2, and individuals with DPRN104 alleles at hl13 always carry Mnas.pTAC14. (C) Location of different copies of pTAC14 based on our mapping experiment.

To examine sequence similarity among *Mimulus pTAC14* genes, we obtained full-length genomic sequences from both DPR parents and generated a neighbor-joining tree (Fig 4, Fig S3). As expected, chromosome 13 copies of *pTAC14* from DPRG102 and IM62 cluster together (*Mg.pTAC14*_*1* and *Migut.M02023*). Likewise, chromosome 14 copies of *pTAC14* from DPRG102 and IM62 cluster together (*Mg.pTAC14*_*2* and *Migut.000467*). However, somewhat counterintuitively, the DPRN104 copy (*Mn.pTAC14*_*1*), which is located on chromosome 13, clusters more closely with *M. guttatus* copies on chromosome 14 than with copies on chromosome 13 (Fig 4B).

**Fig 4:**
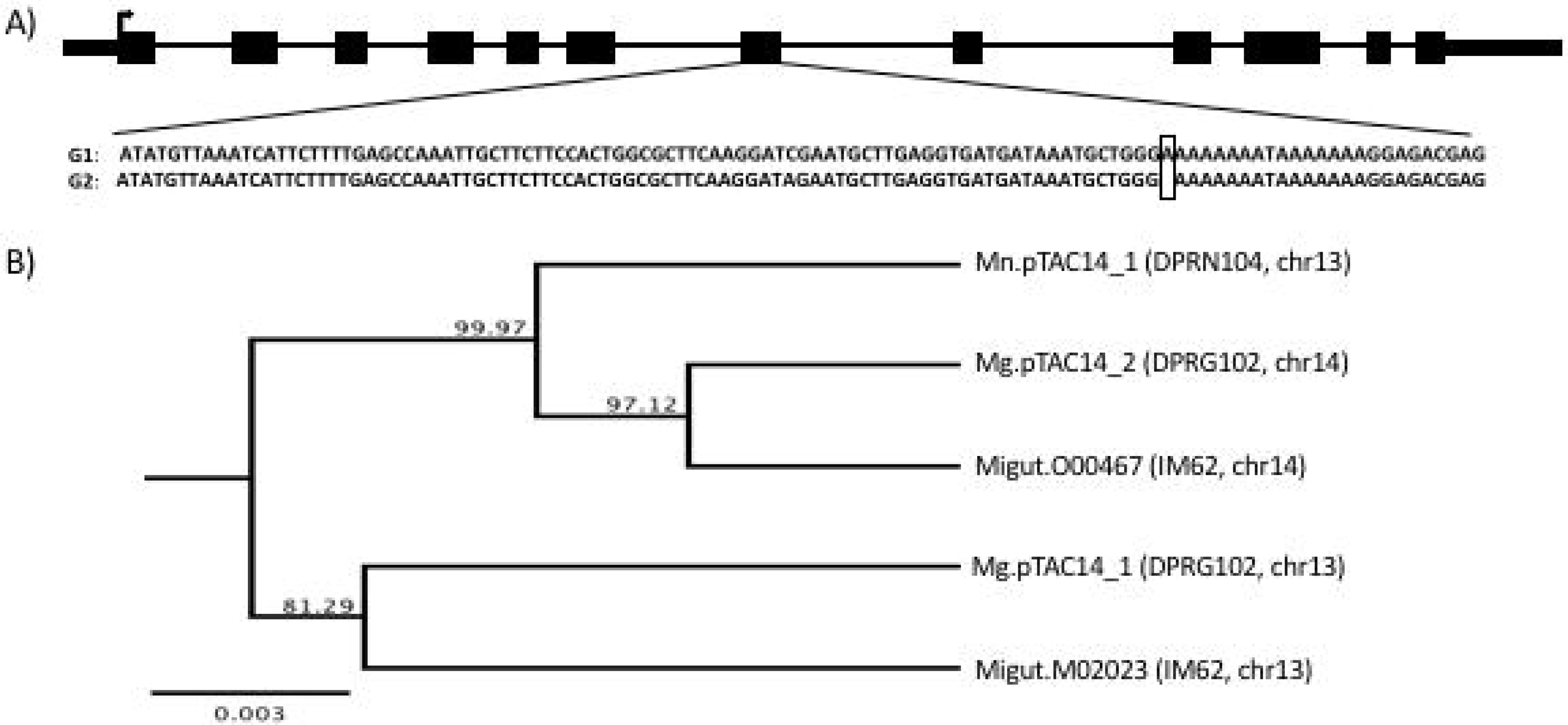
*pTAC14* gene structure and neighbor-joining tree. (A) Gene model of *pTAC14* in *Mimulus* is shown along the top. A frameshift mutation in the G1 copy of *pTAC14* (DPRG102, *Mg.pTAC14_1*) is caused by the insertion of an adenine in the 7^th^ exon, highlighted with a box. (B) Unrooted neighbor-joining tree of *pTAC14* genes from DPRG102, DPRN104, and the IM62 reference genome. Bootstrap support given at node and substitution rate shown for scale.

### *Mimulus pTAC14* duplicates are nonfunctional in hybrid lethal seedlings

Consistent with a causal role for *Mimulus pTAC14* duplicates in hybrid lethality, we discovered several lines of evidence that suggest only one of the two copies is functional in DPRG102. First, *Mg.pTAC14_1* (at *hl13*) contains a frameshift mutation in exon 7, which results in the production of numerous premature stop codons in downstream sequence (Fig 4A, Fig S3). Second, from each inbred line, we PCR-amplified only a single copy of the gene from leaf cDNA: *Mg.pTAC14*_*2* in DPRG102 and *Mn.pTAC14*_*1* in DPRN104. Third, *pTAC14* expression is nearly absent in white F2 hybrid seedlings, which inherit *hl13* from DPRG102 (containing *Mg.pTAC14*_*1*) and *hl14* from DPRN104 (containing no copy of *pTAC14*). Using qPCR and primers that amplify both *Mimulus pTAC14* duplicates, we found strong expression in green parental and F2 seedlings, but not in white F2 seedlings (qPCR on eight additional functional candidates in the *hl13* and *hl14* intervals showed no association between expression and seedling phenotype; Table S2, Fig S4). Additionally, we performed RNAseq on DPRG102, DPRN104, green F2, and white F2 seedlings. Consensus sequences generated from *de novo* assemblies of DPRG102 reads that align to *Migut.M02023* and/or *Migut.000467* (high sequence similarity between *pTAC14* duplicates means that reads align equally well to both copies) correspond to *Mg.pTAC14*_*2*; consensus sequences generated in the same manner from DPRN104 reads correspond to *Mn.pTAC14*_*1*. Moreover, RNAseq SNP variation in green F2 seedlings suggests they express only *Mg.pTAC14*_*2* from DPRG102 and/or *Mn.pTAC14*_*1* from DPRN104. In contrast, read coverage of *pTAC14* transcripts in white F2 seedlings is exceptionally low: of the 1,092 genes that are significantly differentially expressed between white F2 seedlings and green seedlings (DPRG102, DPRN104, green F2 hybrids), the duplicate copies of *pTAC14* are the two most underexpressed (Fig 5). Taken together, these results provide strong evidence that *Mimulus* hybrid lethality is caused by nonfunctional *pTAC14* duplicates: white hybrid seedlings carry unexpressed *Mg.pTAC14*_*1* alleles at *hl13* and are missing *pTAC14* alleles altogether at *hl14*.

**Fig 5:**
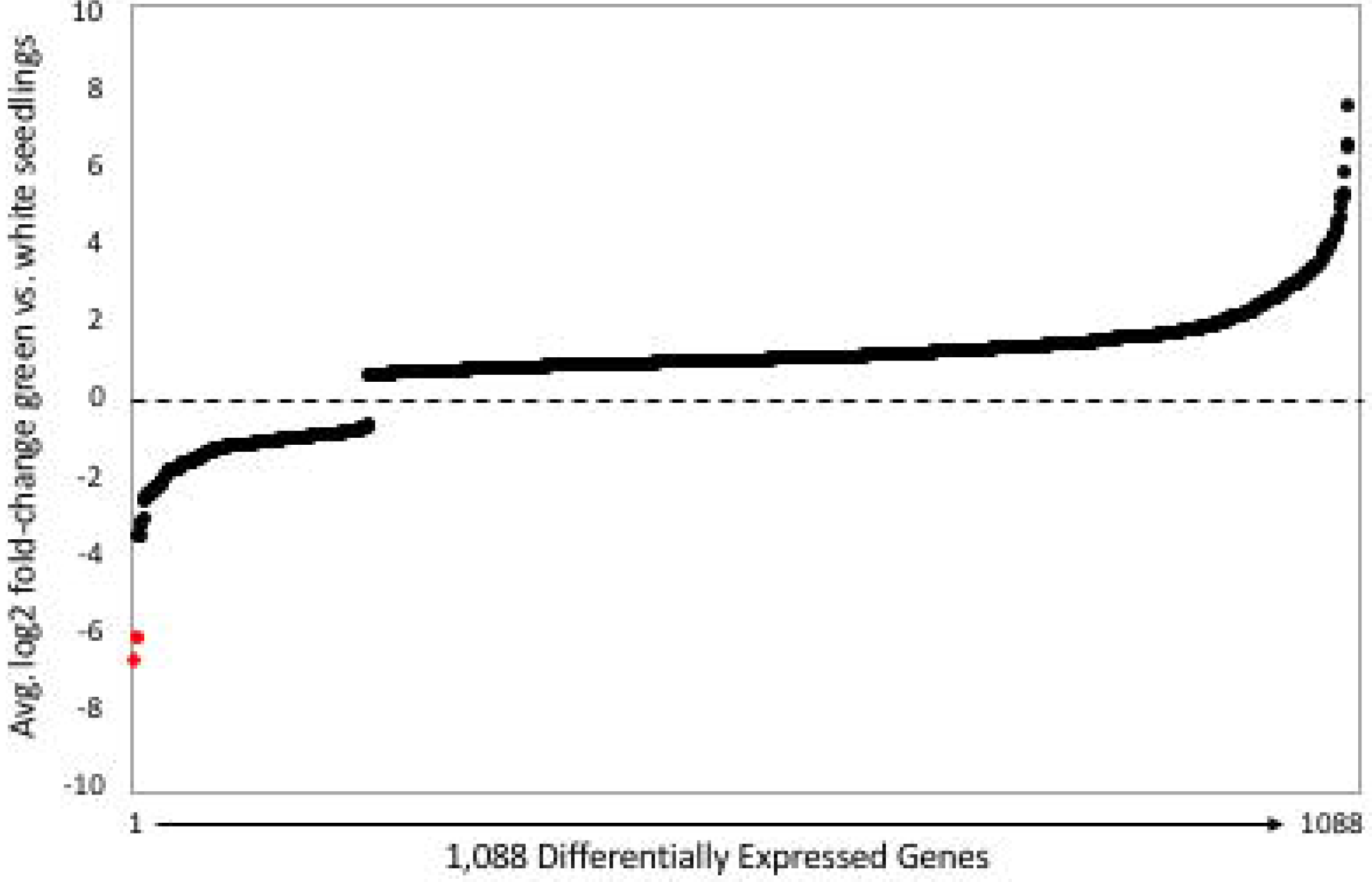
Genome-wide patterns of differential expression. Average log2 fold-change among the 1,092 genes (representing 3% of annotated genes) that are significantly differentially expressed in all pairwise comparisons of white (F2) and green (DPRG102, DPRN104, F2) seedlings. Genes above and below line are over-and underexpressed in white seedlings, respectively. Annotated pTAC14 duplicates are shown as red dots.

### Genome-wide misexpression in hybrid lethal seedlings

Comparison of genome-wide RNAseq patterns among DPRG102, DPRN104, green F2, and white F2 seedlings provides additional support for disrupted *pTAC14* function as a cause of hybrid lethality. White F2 seedlings show a strong signature of genome-wide misexpression: of 27,948 annotated genes, 1,092 (3%) are significantly misexpressed in all three pairwise comparisons between white and green seedlings (Fig 5). Among transcripts that are underexpressed in white seedlings (*N* = 209), we found a significant enrichment of genes involved in photosynthesis and/or located within the thylakoid and photosynthetic membranes. Among overexpressed transcripts (*N* = 883), we observed an enrichment of heat shock proteins and glutathione peroxidase proteins (Table S3). Furthermore, consistent with disrupted *pTAC14* function, we discovered evidence for severe misexpression of chloroplast-encoded genes in white seedlings (Fig 6, Fig S5). In *A. thaliana*, knockouts of *pTAC14* disable the PEP (plastid-encoded bacterial type) RNA polymerase, which leads to reduced transcription of some chloroplast-encoded genes, particularly those involved in photosynthesis (*e.g*., photosystem I, photosystem II, and cytochrome b6f), and increased transcription of others such as the *rpo* genes (Gao et al. 2011). Of 52 putative *Mimulus* chloroplast genes, those involved in photosynthesis, and thus likely to be transcribed by PEP RNA polymerase, were often significantly underexpressed in white F2 seedlings. In contrast, homologs of *A. thaliana rpo* genes were significantly overexpressed. Several additional putative chloroplast genes (*e.g*., ATP synthase, NADH Dehydrogenase, ribosomal proteins) that are likely transcribed by both PEP and the nuclear-encoded phage-type (NEP) RNA polymerase [44, 45] were also significantly misexpressed (both up- and downregulated) in white seedlings. Taken together, these patterns of gene misexpression in *Mimulus* F2 white seedlings, which show a remarkable similarity to patterns observed in *A. thaliana pTAC14* knockouts [42], provide strong evidence for a causal role of *pTAC14* duplicates in *Mimulus* hybrid lethality.

**Fig 6:**
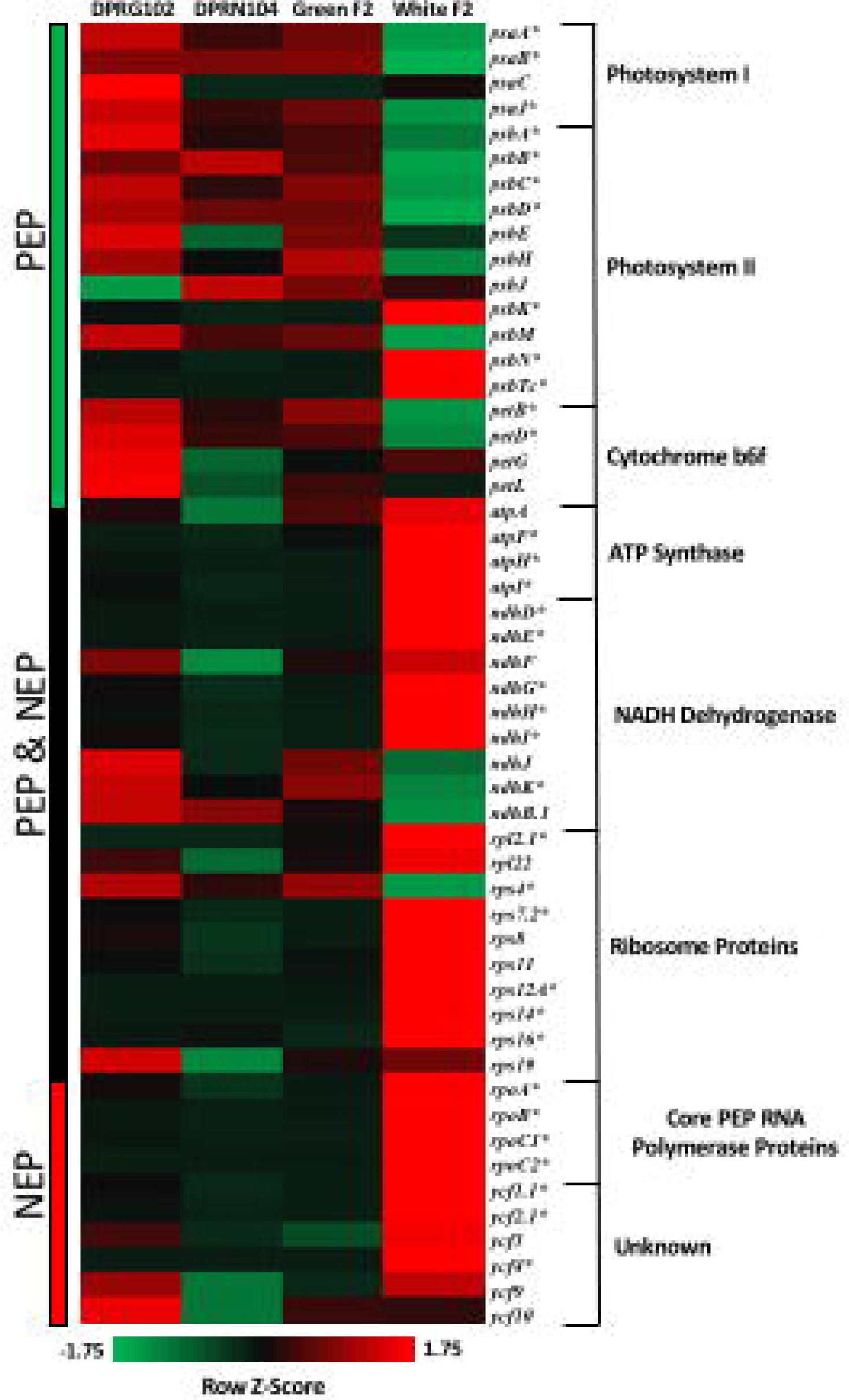
Misexpression of chloroplast genome in white seedlings. Heat map displaying expression patterns of 52 chloroplast genes from seven chloroplast gene families in seedlings from DPRG102, DPRN104, Green F2s and White F2s. Z-scores of normalized FPKM values were calculated for each row to illustrate relative expression differences among genes. Bars on left of heatmap indicate whether genes are primarily transcribed by PEP, PEP and NEP, or NEP RNA polymerases. *denotes genes that are significantly differentially expressed between white seedlings and all green seedlings (DPRG102, DPRN104, and Green F2s).

## Discussion

Identifying the molecular genetic basis of hybrid incompatibilities between recently diverged, wild species is a critical first step toward understanding their evolutionary origins and role in speciation. We have shown that duplicate copies of *Mimulus pTAC14*, a gene critical for chloroplast development in *A. thaliana* [42], causes hybrid lethality between sympatric *M. guttatus* and *M. nasutus* at the DPR site. We fine-mapped hybrid lethality to *hl13* and *hl14*, two small nuclear genomic regions on chromosomes 13 and 14. In DPRG102 (*M. guttatus*), *pTAC14* is present in each of these genomic intervals, but only the *hl14* copy is expressed. In DPRN104 (*M. nasutus*), *pTAC14* is present only in the *hl13* interval, consistent with either of two possibilities: the *hl13* copy is ancestral and this line lacks the duplication, or a large deletion has removed all trace of the gene from *hl14*. As a consequence of divergent resolution of these duplicate genes, F2 hybrids that are homozygous for DPRG102 alleles at *hl13* and homozygous for DPRN102 alleles at *hl14* contain no functional copy of *Mimulus pTAC14*. These hybrids fail to produce chlorophyll and die in the cotyledon stage of development, remarkably similar to what is observed in *pTAC14* knockouts in *A. thaliana* [42]. To our knowledge, this is the first pair of hybrid incompatibility genes identified between naturally hybridizing species.

Using complementary genetic mapping and functional genomics approaches, our study provides strong evidence that nonfunctional *Mimulus pTAC14* is the cause of DPRG102-DPRN104 hybrid lethality. In *A. thaliana, pTAC14* is one of several nuclear-encoded proteins that are critical components of the PEP RNA polymerase. As the only RNA polymerase responsible for transcribing key plastid-encoded photosynthesis genes (photosystem I, photosystem II, cytochrome b6f), PEP is an essential enzyme in plants [44, 46]. Knockouts that disrupt or inactivate PEP activity (such as *pTAC14*) share several common phenotypes, including the complete lack of photosynthesis, down-regulation of plastid-encoded photosynthesis genes, and up-regulation of plastid-encoded PEP subunits (e.g., *rpo* genes) [42, 45-50]. The transcriptional profile of putative chloroplast genes in white DPRN104-DPRG102 F2 seedlings bears a striking resemblance to that of *A. thaliana* mutants that disable PEP, particularly the down-regulation of photosynthetic genes and up-regulation of *rpo* genes. This fact, combined with our finding that *pTAC14* is the only PEP-associated protein that maps to *hl13* or *hl14* provides strong evidence that hybrid lethality is the product of PEP-inactivation

But what causes the lack of *pTAC14* expression in white hybrid seedlings? In *M. nasutus* (DPRN104), because *pTAC14* is missing entirely from the *hl14* interval, the *hl13* copy (*Mn.pTAC14_1*) is the only one expressed. In *M. guttatus* (DPRG102), the situation is less clear. Although both copies (*Mg.pTAC14_1* at *hl13* and *Mg.pTAC14_2* at *hl14*) are present and highly similar in exons (Fig S3), our qPCR and RNAseq experiments demonstrate that only one of them – *Mg.pTAC14_2* – is expressed. Further work will be required to determine the molecular nature of this change in gene expression. The most obvious possibility is that non-sense mediated decay has efficiently targeted *Mg.pTAC14_1*, which carries a series of premature stop codons. Another possibility, is that a cis-regulatory mutation disrupts *Mg.pTAC14_1* transcription in DPRG102. Alternatively, expression might be prevented by the epigenetic silencing of one duplicate by the other, as was recently shown for sterile and lethal combinations segregating within *A. thaliana* [28, 51]. Whatever its cause, disrupted expression is not the only problem with DPRG102 *Mg.pTAC14_1;* this gene copy also carries a 1-bp insertion that, if transcribed, would result in a truncated, and potentially nonfunctional, protein. We do not yet know which of these two functional changes to DPRG102 *Mg.pTAC14_1* arose first.

The evolution of hybrid lethality in this system thus appears entirely consistent with a scenario of duplication and neutral non-functionalization within *M. guttatus*. Given the ubiquity of gene duplications in plant and animal genomes, divergent resolution of paralogs due to degenerative mutation and genetic drift has been proposed as a major source of hybrid incompatibilities [24, 25, 52]. Although initially redundant duplicate genes might sometimes evolve new or partial functions favored by selection [53], our study and others suggest that duplicates involved in hybrid incompatibilities are more often subject to mutations that disable function in one copy. Within *A. thaliana* and between closely related *Oryza* species, divergent resolution of duplicates has occurred through nonsense mutations [20, 22] and disruptions to expression [21, 27, 51]. In a more distantly related species pair of *Drosophila*, hybrid sterility is caused by a gene transposition, with degenerative mutations having presumably removed any remnant of the duplication that likely preceded its evolution [26]. Remarkably, then, to explain the evolution of hybrid dysfunction in *Mimulus* and several other diverse systems, there is no need to invoke processes beyond mutation and genetic drift.

In addition to showing that *Mimulus* hybrid lethality is due to nonfunctional *pTAC14*, our analyses have begun to provide some insight into the duplication history of this gene. As might be expected, within *M. guttatus* (DPRG102 and IM62), *pTAC14* copies on chromosome 13 are most related and *pTAC14* copies on chromosome 14 are most related (Fig 4). However, somewhat counterintuitively, *pTAC14* from DPRN104, which is located on chromosome 13, is most closely related to the *M. guttatus* copies on chromosome 14. We interpret this finding, along with the fact that we find no trace of *pTAC14_2* at *hl14* in DPRN104, as evidence that *Mimulus pTAC14_1* on chromosome 13 is the ancestral copy. Under this scenario, both the duplicate copy on chromosome 14 (*Mg.pTAC14_2*) and the *M. nasutus* copy on chromosome 13 (*Mn.pTAC14_1*) would have arisen from a similar genetic variant (Fig 7). Standing genetic variation within and between populations of *M. guttatus* is high [29, 54-56, 57] so it is likely that ancestral populations carried multiple variants of *pTAC14_1*. Both the duplicated copy in *M. guttatus* (*Mg.pTAC14_2*) and the ancestral copy in the selfing *M. nasutus* (*Mn.pTAC14_1*) would be expected to carry only a small subset of ancestral variation. Unfortunately, we have not yet been able to assess molecular patterns of *Mimulus pTAC14* variation in a wider sample of *M. guttatus* and *M. nasutus*. Although whole genome resequence data are available from a number of lines [29, 54, 57], short-read sequences of *Mimulus pTAC14* align equally well to both annotated copies in the IM62 reference genome (*Migut.M02023* and *Migut.000467*). Once *Mimulus pTAC14* is sequenced from a broader sample of individuals, we speculate that *Mn.pTAC14_1* from *M. nasutus* and *Mg.pTAC14_2* from *M. guttatus* will cluster as distinct monophyletic groups nested within the greater diversity of sequences present at the ancestral *Mg.pTAC14_1* from *M. guttatus*. Interestingly, white seedlings are often observed segregating at low frequencies within *M. guttatus* populations, which manifest as epistatic inbreeding depression [58, 59] that may be due to divergent resolution of duplicate genes similar to the one characterized here. Indeed, variation for functional and non-functional *pTAC14* variations exists at both *hl13* and *hl14* in *M. guttatus*, indicating that this duplication may present such a case (Zuellig and Sweigart, unpublished results).

**Figure 7:**
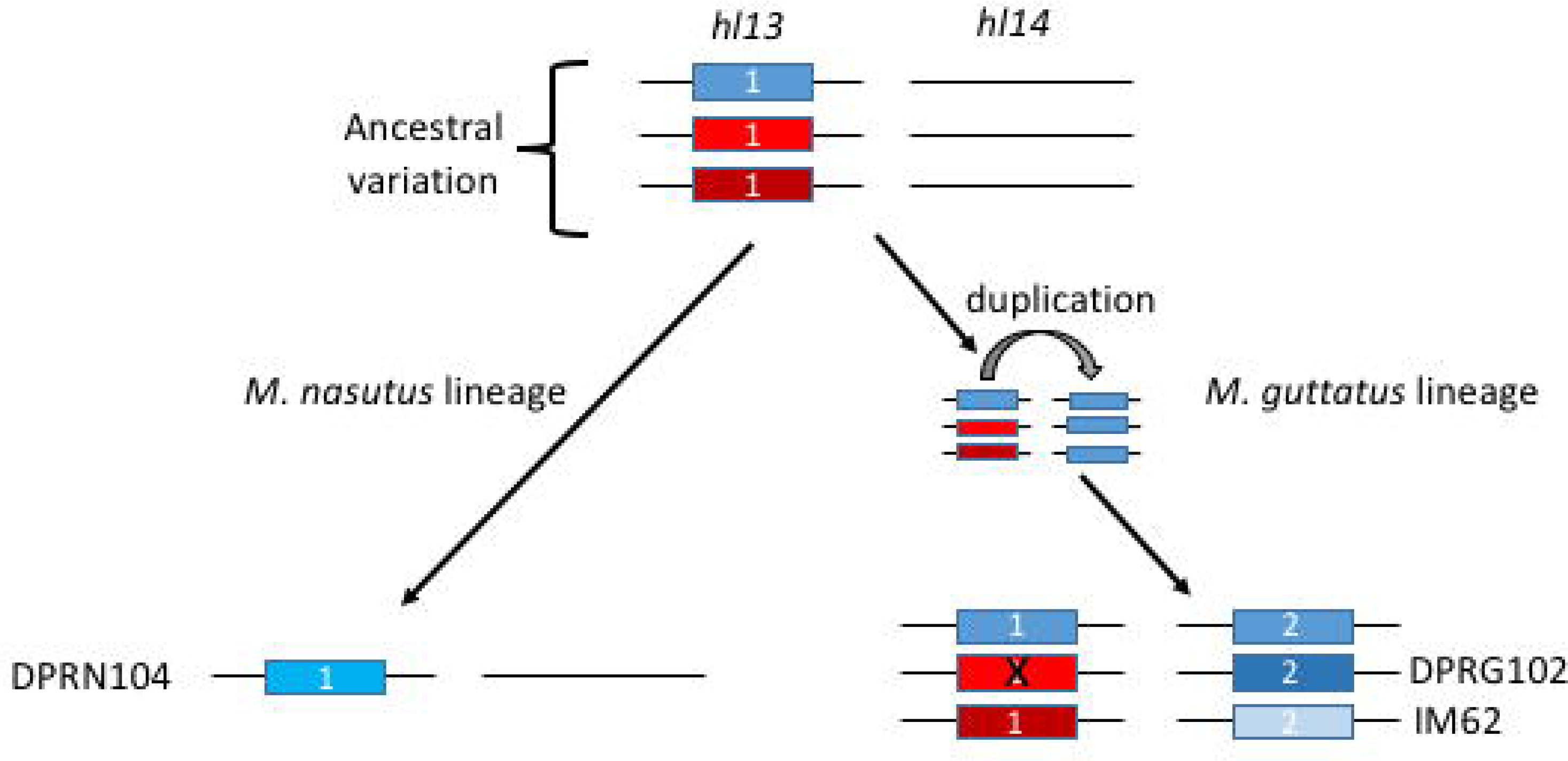
Hypothetical model for the evolution of *pTAC14*. Prior to gene duplication the ancestral *M. guttatus*-like population harbored genetic variation for *pTAC14* at *hl13* (different colored boxes represent genetic variation at *pTAC14*). During speciation, *M. nasutus* acquired a copy of *pTAC14* harboring ‘blue’ variation (left). Similarly, the *M. guttatus*-specific duplication involved a closely related ‘blue’ *pTAC14* variant. Contemporary variation within *M. nasutus* and *pTAC14* variants located at *hl14* resemble one another genetically due to common ancestry, while *pTAC14* variants at *hl13* in *M. guttatus* continue to harbor considerable genetic variation, including variants that contain functional (IM62) and non-functional (DPRG102) variants.

Our study provides the first detailed study of hybrid incompatibility genes from naturally hybridizing species and contributes to a growing body of literature that shows hybrid incompatibilities can result from neutral evolutionary change within species. Going forward, it will be important to address whether these barriers can persist in the face of ongoing gene flow. Theoretical treatments of this question have consistently concluded that the maintenance of hybrid incompatibility alleles between hybridizing populations relies heavily on a selective advantage within species [60–64]. If so, neutrally evolving hybrid incompatibility alleles might be precluded from affecting reproductive isolation in any more than a transient fashion, with gene flow temporarily constrained until the hybrid incompatibility degrades with time. Nevertheless, other factors such as strong linkage to selected alleles (e.g., [16]) and constraints on gene dosage (e.g., [65]) may play an important role in such incompatibilities. By showing that the duplication of *pTAC14* underlies hybrid lethality among sympatric *Mimulus* species, we now have a natural system in place to test broader questions regarding the evolutionary significance of neutral processes on speciation.

## Materials and Methods

### *Mimulus* lines and genetic crosses

We generated inbred lines of *M. guttatus* and *M. nasutus* derived from wild individuals collected from Don Pedro Reservoir (DPR) in central California [33]. Wild-collected seed was sown on moist Fafard 3-B potting soil in 2.5” pots, cold-stratified in the dark at 4C for two weeks, and moved the UGA greenhouses to germinate under 16 hour days at 23°C (growth conditions were constant for all experiment, though RNAseq experiment took place in growth chamber). Upon germination, a single seedling was transplanted to its own 2.5” pot, allowed to flower, and self-fertilized. After three generations of selfing with single-seed descent, each line (DPRG102: *M. guttatus*; DPRN104: *M. nasutus*) was intercrossed to generate reciprocal F1 and F2 hybrids (maternal parent always listed first in crosses).

### Molecular analyses and whole genome sequencing

DNA was extracted from seedlings and adult leaf tissue using a standard CTAB-chloroform protocol [66] modified for use in a 96-well format. Genotyping was performed using a combination of exon-primed intron-spanning size polymorphic markers containing 5’ fluorescent tags (6-FAM or HEX) and SNP-containing gene fragments that were analyzed through Sanger sequencing. We designed size-polymorphic and SNP markers from polymorphisms observed in whole genome re-sequence data of multiple lines of *M. guttatus* and *M. nasutus* [29, 67, 68] and confirmed polymorphisms by genotyping parental lines used in our study. A standard touchdown PCR protocol was used in all amplifications and Sanger sequencing reactions were prepared using BigDye v3.1 mastermix (Applied Biosystems, Foster City, USA). Genotyping and Sanger sequencing reactions were run on an ABI3730XL automated DNA sequencer at the Georgia Genomics Facility and analyzed using GENEMARKER [69] and Sequencher (Gene Codes Corporation, Ann Arbor, USA) software, respectively.

For whole genome sequencing of bulked segregants, we generated equimolar amounts of DNA from green (N=26) and white hybrid seedlings (N=34). Green and white DNA was pooled separately and sent to the Duke Center for Genomic and Computational Biology, where Illumina libraries with unique barcodes were prepared and sequenced using the Illumina Hi-seq platform (100bp single-end reads). Reads from both pools were aligned to the *M. guttatus* (IM62) reference genome (https://phytozome.jgi.doe.gov), along with previously generated whole genome re-sequence data for DPRN104 [29]. Reads were aligned using Burrows-Wheeler Aligner (bwa, [70]) with a minimum alignment quality threshold of Q29 (filtering done with samtools, [71]). We identified 235,922 SNPs that differentiated the IM62 reference genome from DPRN104 using the samtools mpileup function, which provided a list of SNPs that differentiate these two lineages. We used the samtools mpileup function to estimate the frequency of each SNP (‘alternate allele frequency’) within white and green BSA pools. Since SNPs were not based on differences between DPRN104 and DPRG102, our analysis assumes that *M. guttatus* lines IM62 and DPRG102 (which is not sequenced) share a common set of SNPs.

### qPCR and RNA sequencing

We performed quantitative PCR on a subset of strong candidate genes within *hl13* and *hl14* (9 genes total, Table S2), comparing expression patterns in seedlings from DPRG102, DPRN104, green F2s, and white F2s. We extracted RNA from pools of 10 seedlings for each genotypic class using a Zymo MicroRNA Kit (Zymo Research, Irvine, USA) followed by cDNA synthesis with GOscript Reverse Transcriptase (Promega, Madison, USA). We designed exon-specific primers to amplify fragments of each gene (Table S4), amplified fragments using standard touchdown PCR, and visualized gene fragments on a 1% agarose gel.

We performed an RNAseq experiment to compare genome-wide expression profiles between white and green seedlings. We used lines of DPRG102 and DPRN104 that had been inbred for 5 generations and their green and white F2 progeny, which resulted in three classes of green seedlings (DPRG102, DPRN104, and green F2s) and a single class of white seedlings (white F2s). Seedlings with fully expanded cotyledons began to emerge within 3 days and continued to emerge for a week thereafter. We collected pools of 10 seedlings from each biological class directly into 2mL Eppendorf tubes filled with liquid nitrogen. We then extracted RNA from these pools using the Zymo Quick-RNA microprep kit (Zymo Research, Irvine, USA) and estimated RNA concentration using a qubit fluorometer (Life Technologies, Paisley, UK). High quality RNA was subsequently submitted to the Duke Center for Genomic and Computational Biology, where Kapa Stranded mRNA-Seq libraries (Kapa Biosystems, Wilmington, USA) were prepared and samples were sequenced across a single lane of Illumina Hiseq 4000 with single-end 50 bp reads. In total, our analysis involved three replicates each of DPRG102 and DPR104 green seedlings, five replicates of green F2 seedlings, and six replicates of white F2 seedlings, where each replicate was a pool of 10 seedlings.

We utilized the cufflinks pipeline [72] to assess patterns of differential expression among the four genotypic classes (DPRG102, DPRN104, green F2s, and white F2s). We aligned trimmed and filtered reads (Q>20) to the *M. guttatus* IM62 reference genome in TopHat2 [73], which resulted in an average of 19 million reads aligned per biological replicate. We then assembled transcriptomes in cufflinks, using the IM62 reference transcriptome as a guide. We used ‘cuffnorm’ to normalize transcript abundance for each genotypic class and ‘cuffdiff’ to calculate differential expression for all pairwise comparisons. For data management and sorting, we used Microsoft excel, the R statistical package [74], and the R package CummeRbund [72]. Gene ontology (GO) enrichment analyses were carried out for particular subsets of data that exhibited patterns of differential expression between white and green seedlings. To perform these analyses, we used GOstat [75] and GO::TermFinder [76] implemented in the Phytomine user interface (https://phytozome.jgi.doe.gov). For GO term analyses, we used all annotated genes in the v2.0 IM62 *M. guttatus* reference assembly to serve as the background population and used a Bonferroni cutoff value of 0.05 to test for significant GO term enrichment in our subset of differentially expressed genes. For our analysis of chloroplast-encoded genes, we generated a list of putative chloroplast genes, since no chloroplast genome assembly is currently available for *M. guttatus*. We generated this set by first downloading a list of 135 genes present in the chloroplast genome in *A. thaliana* from the TAIR database (www.arabidopsis.org). We used this list to identify *M. guttatus* homologs in the Phytomine database (https://phytozome.jgi.doe.gov). This approach yielded a set of 52 putative *Mimulus* chloroplast genes that are currently included (and, presumably, misassembled) in the nuclear genome (no homologs were identified for the other 83 genes used in our search). A substantial fraction of these genes (69%) occur along chromosome 4 from positions 6,719,000-7,985,375, which contains 183 genes total.

### Gene sequencing and phylogenetic analyses

To obtain full-length *pTAC14* sequences from DPRG102 and DPRN104, we amplified both genomic and cDNA using primers designed within conserved exonic sequence. PCR fragments were amplified using Phusion High-Fidelity DNA Polymerase (New England Biolabs, Ipswich, USA) and either directly sequenced or sequenced after cloning into the TOPO TA Vector (Thermo Fisher, Carlsbad, USA). Additionally, we extracted full-length transcript sequences from RNAseq data by performing *de novo* assemblies on reads that mapped to candidate genes using the Geneious Assembler (Biomatters, Newark, USA). When reads mapped to duplicated genes, they were combined into a single *de novo* assembly and 95% confidence consensus sequences were constructed. Using PHYML [77], we constructed a neighbor-joining tree for *pTAC14* using 4,319 bp of genomic sequence (excluding 5’ and 3’ UTRs and insertions/deletions coded as single variants) with branch support determined with 1000 bootstraps. For the tree presented in Fig 4b, we used the general time-reversible model with four substitution rate categories and allowed the program to estimate the proportion of variable sites and the gamma distribution parameter (varying these parameters produced identical consensus trees).

## Acknowledgments

We thank Noland Martin and John Willis for first identifying white seedlings at Don Pedro Reservoir and encouraging us to study them. We thank Amanda Kenney for her help with BSA experiments and intellectual contributions. John Willis, Nick Arthur, Rachel Kerwin, Nick Batora, and Sam Mantel provided thoughtful comments, which significantly improved the quality of our manuscript.

---

**Fig S1: Hybrid lethality phenotype.** White seedlings segregate in 1:15 in reciprocal F2 hybrids of *M. guttatus* (DPRG102) and *M. nasutus* (DPRN104). Photos kindly provided by Adam J Bewick.

**Fig S2: Bulked Segregant Analysis of green and white F3 pools.** Difference in average allele frequency between green and white pools (plotted along the fourteen *Mimulus* chromosomes) was calculated in 200-SNP windows with 100-SNP overlap. The 0.5% most divergent windows are highlighted as black dots, which are all located at the distal end of chromosome 13 in contiguous windows and represent the candidate *hl13* region. Red dots, which overlap with the previously mapped interval for *hl14*, show the 0.5% least divergent windows (calculated as absolute difference in allele frequency).

**Fig S3: Genomic sequence alignment of *pTAC14* variants.** Exons are highlighted blue. Black boxes show ten SNPs that were used to positionally map copies *Mg.pTAC14_1, Mg.pTAC14_2, and Mn.pTAC14_1* in DPRN104 x DPRG102 F2 seedlings. Red box indicates site of frameshift mutation in *Mg.pTAC14_1*.

**Fig S4: *pTAC14* is not expressed in white seedlings.** PCR products run on 1% agarose gel with 2-log ladder. Control transcript is Migut.M00195 (ACYL-COENZYME A OXIDASE-LIKE PROTEIN). Note that DNA and RNA was extracted from pools of 10 seedlings for each genotype.

**Fig S5: Misexpression of key photosynthetic genes indicates PEP inactivity.** Differential expression among chloroplast genes transcribed by PEP (*psaA, psaB, psbA, psbB, petB, petD*), PEP and NEP (*atpA* and *ndhB*), and NEP (*rpoA*, *rpoB*, and *rpoC1*). Green: average log2 fold-change in all pairwise comparisons among green seedlings (DPRG102, DPRN104, Green F2). White: average log2 fold-change in pairwise comparisons between white F2 seedlings and green seedlings. *Significantly down-regulated in white seedlings (all pairwise comparisons, p<5.0^-5^). **Significantly up-regulated in white seedlings (all pairwise comparisons, p<5.0^-5^).

**Table S1:** Segregation of white seedlings in parental and reciprocal hybrid crosses.

**Table S2:** Candidate genes at *hl13* and *hl14*.

**Table S3:** GO term enrichment of differentially expressed genes

**Table S4:** Genotyping and sequencing primers used for fine-mapping.

## References

1. Bateson W. Heredity and variation in modern lights. Darwin and modern science. 1909;85:101.

2. Dobzhansky T, Dobzhansky TG. Genetics and the Origin of Species: Columbia University Press; 1937.

3. Muller HJ, editor Isolating mechanisms, evolution and temperature. Biol Symp; 1942.

4. Fishman L, Sweigart AL. When two rights make a wrong: evolutionary genetics of plant hybrid incompatibilities. Annual review of plant biology. in press.

5. Maheshwari S, Barbash DA. The Genetics of Hybrid Incompatibilities. Annual Review of Genetics 2011;45(1):331–55. doi:10.1146/annurev-genet-110410-132514. PubMed PMID: 21910629.

6. Presgraves DC. The molecular evolutionary basis of species formation. Nature Reviews Genetics. 2010;11(3):175–80.

7. Sweigart AL, Willis JH. Molecular evolution and genetics of postzygotic reproductive isolation in plants. F1000 biology reports. 2012;4:23.

8. Brideau NJ, Flores HA, Wang J, Maheshwari S, Wang X, Barbash DA. Two Dobzhansky-Muller Genes Interact to Cause Hybrid Lethality in Drosophila. Science. 2006;314(5803):1292–5. doi:10.1126/science.1133953.

9. Maheshwari S, Wang J, Barbash DA. Recurrent Positive Selection of the Drosophila Hybrid Incompatibility Gene Hmr. Molecular Biology and Evolution. 2008;25(11):2421–30. doi:10.1093/molbev/msn190.

10. Oliver PL, Goodstadt L, Bayes JJ, Birtle Z, Roach KC, Phadnis N, et al. Accelerated evolution of the Prdm9 speciation gene across diverse metazoan taxa. PLoS Genet. 2009;5(12):e1000753.

11. Phadnis N, Orr HA. A single gene causes both male sterility and segregation distortion in Drosophila hybrids. science. 2009;323(5912):376–9.

12. Presgraves DC, Balagopalan L, Abmayr SM, Orr HA. Adaptive evolution drives divergence of a hybrid inviability gene between two species of Drosophila. Nature. 2003;423(6941):715–9.

13. Tang S, Presgraves DC. Evolution of the Drosophila nuclear pore complex results in multiple hybrid incompatibilities. Science. 2009;323(5915):779–82.

14. Chen C, Chen H, Lin Y-S, Shen J-B, Shan J-X, Qi P, et al. A two-locus interaction causes interspecific hybrid weakness in rice. Nature communications. 2014;5:3357.

15. Sicard A, Kappel C, Josephs EB, Lee YW, Marona C, Stinchcombe JR, et al. Divergent sorting of a balanced ancestral polymorphism underlies the establishment of gene-flow barriers in Capsella. Nature Communications. 2015;6:7960. doi:10.1038/ncomms8960 http://www.nature.com/articles/ncomms8960-supplementary-information.

16. Wright KM, Lloyd D, Lowry DB, Macnair MR, Willis JH. Indirect evolution of hybrid lethality due to linkage with selected locus in Mimulus guttatus. PLoS Biol. 2013;11(2):e1001497.

17. Case AL, Finseth FR, Barr CM, Fishman L. Selfish evolution of cytonuclear hybrid incompatibility in &<em&>Mimulus&</em&>. Proceedings of the Royal Society B: Biological Sciences. 2016;283(1838).

18. Tao Y, Hartl DL, Laurie CC. Sex-ratio segregation distortion associated with reproductive isolation in Drosophila. Proceedings of the National Academy of Sciences. 2001;98(23):13183–8. doi:10.1073/pnas.231478798.

19. Zhang L, Sun T, Woldesellassie F, Xiao H, Tao Y. Sex Ratio Meiotic Drive as a Plausible Evolutionary Mechanism for Hybrid Male Sterility. PLOS Genetics. 2015;11(3):e1005073. doi:10.1371/journal.pgen.1005073.

20. Mizuta Y, Harushima Y, Kurata N. Rice pollen hybrid incompatibility caused by reciprocal gene loss of duplicated genes. Proceedings of the National Academy of Sciences. 2010;107(47):20417–22. doi:10.1073/pnas.1003124107.

21. Nguyen GN, Yamagata Y, Shigematsu Y, Watanabe M, Miyazaki Y, Doi K, et al. Duplication and Loss of Function of Genes Encoding RNA Polymerase III Subunit C4 Causes Hybrid Incompatibility in Rice. G3: Genes|Genomes|Genetics. 2017;7(8):2565.

22. Yamagata Y, Yamamoto E, Aya K, Win KT, Doi K, Sobrizal, et al. Mitochondrial gene in the nuclear genome induces reproductive barrier in rice. Proceedings of the National Academy of Sciences. 2010;107(4):1494–9. doi:10.1073/pnas.0908283107.

23. Oka H-I. Analysis of Genes Controlling F(1) Sterility in Rice by the Use of Isogenic Lines. Genetics. 1974;77(3):521–34. PubMed PMID: PMC1213144.

24. Lynch M, Force AG. The Origin of Interspecific Genomic Incompatibility via Gene Duplication. The American Naturalist. 2000;156(6):590–605. doi: doi:10.1086/316992.

25. Werth CR, Windham MD. A Model for Divergent, Allopatric Speciation of Polyploid Pteridophytes Resulting from Silencing of Duplicate-Gene Expression. The American Naturalist. 1991;137(4):515–26. doi: doi:10.1086/285180.

26. Masly JP, Jones CD, Noor MA, Locke J, Orr HA. Gene transposition as a cause of hybrid sterility in Drosophila. Science. 2006;313(5792):1448–50.

27. Bikard D, Patel D, Le Metté C, Giorgi V, Camilleri C, Bennett MJ, et al. Divergent Evolution of Duplicate Genes Leads to Genetic Incompatibilities Within A. thaliana. Science. 2009;323(5914):623–6. doi:10.1126/science.1165917.

28. Durand S, Bouché N, Strand EP, Loudet O, Camilleri C. Rapid establishment of genetic incompatibility through natural epigenetic variation. Current Biology. 2012;22(4):326–31.

29. Brandvain Y, Kenney AM, Flagel L, Coop G, Sweigart AL. Speciation and introgression between Mimulus nasutus and Mimulus guttatus. PLoS Genet. 2014;10(6):e1004410.

30. Diaz A, Macnair M. Pollen tube competition as a mechanism of prezygotic reproductive isolation between Mimulus nasutus and its presumed progenitor M. guttatus. New Phytologist. 1999;144(3):471–8.

31. Kenney AM, Sweigart AL. Reproductive isolation and introgression between sympatric Mimulus species. Molecular ecology. 2016;25(11):2499–517.

32. Kiang Y, Hamrick J. Reproductive isolation in the Mimulus guttatus M. nasutus complex. American Midland Naturalist. 1978:269–76.

33. Martin NH, Willis JH. Ecological divergence associated with mating system causes nearly complete reproductive isolation between sympatric Mimulus species. Evolution. 2007;61(1):68–82.

34. Case AL, Willis JH. Hybrid male sterility in Mimulus (Phrymaceae) is associated with a geographically restricted mitochondrial rearrangement. Evolution. 2008;62(5):1026–39.

35. Fishman L, Willis JH. A cytonuclear incompatibility causes anther sterility in Mimulus hybrids. Evolution. 2006;60(7):1372–81.

36. Kiang Y, Libby W. Maintenance of a lethal in a natural population of Mimulus guttatus. The American Naturalist. 1972;106(949):351–67.

37. Martin NH, Willis JH. Geographical variation in postzygotic isolation and its genetic basis within and between two Mimulus species. Philosophical Transactions of the Royal Society of London B: Biological Sciences. 2010;365(1552):2469–78.

38. Sweigart AL, Fishman L, Willis JH. A Simple Genetic Incompatibility Causes Hybrid Male Sterility in Mimulus. Genetics. 2006;172(4):2465–79. doi:10.1534/genetics.105.053686.

39. Sweigart AL, Flagel LE. Evidence of natural selection acting on a polymorphic hybrid incompatibility locus in Mimulus. Genetics. 2015;199(2):543–54.

40. Vickery Jr RK. Case studies in the evolution of species complexes in Mimulus. Evolutionary biology: Springer; 1978. p. 405–507.

41. Sweigart A, Willis J. Patterns of nucleotide diversity are affected by mating system and asymmetric introgression in two species of Mimulus. Evolution. 2003;57:2490–506.

42. Gao Z-P, Yu Q-B, Zhao T-T, Ma Q, Chen G-X, Yang Z-N. A functional component of the transcriptionally active chromosome complex, Arabidopsis pTAC14, interacts with pTAC12/HEMERA and regulates plastid gene expression. Plant physiology. 2011;157(4):1733–45.

43. Hellsten U, Wright KM, Jenkins J, Shu S, Yuan Y, Wessler SR, et al. Fine-scale variation in meiotic recombination in Mimulus inferred from population shotgun sequencing. Proceedings of the National Academy of Sciences. 2013;110(48):19478–82.

44. Hajdukiewicz PT, Allison LA, Maliga P. The two RNA polymerases encoded by the nuclear and the plastid compartments transcribe distinct groups of genes in tobacco plastids. The EMBO journal. 1997;16(13):4041–8.

45. Swiatecka-Hagenbruch M, Liere K, Börner T. High diversity of plastidial promoters in Arabidopsis thaliana. Molecular Genetics and Genomics. 2007;277(6):725–34.

46. Kremnev D, Strand Å. Plastid encoded RNA polymerase activity and expression of photosynthesis genes required for embryo and seed development in Arabidopsis. Frontiers in plant science. 2014;5.

47. Demarsy E, Buhr F, Lambert E, Lerbs-Mache S. Characterization of the plastid-specific germination and seedling establishment transcriptional programme. Journal of experimental botany. 2012;63(2):925–39.

48. Ishizaki Y, Tsunoyama Y, Hatano K, Ando K, Kato K, Shinmyo A, et al. A nuclear-encoded sigma factor, Arabidopsis SIG6, recognizes sigma-70 type chloroplast promoters and regulates early chloroplast development in cotyledons. The Plant Journal. 2005;42(2):133–44.

49. Liere K, Börner T. Transcription and transcriptional regulation in plastids. Cell and molecular biology of plastids: Springer; 2007. p. 121–74.

50. Pfannschmidt T, Blanvillain R, Merendino L, Courtois F, Chevalier F, Liebers M, et al. Plastid RNA polymerases: orchestration of enzymes with different evolutionary origins controls chloroplast biogenesis during the plant life cycle. Journal of experimental botany. 2015;66(22):6957–73.

51. Blevins T, Wang J, Pflieger D, Pontvianne F, Pikaard CS. Hybrid incompatibility caused by an epiallele. Proceedings of the National Academy of Sciences. 2017:201700368.

52. Lynch M, Conery JS. The Evolutionary Fate and Consequences of Duplicate Genes. Science. 2000;290(5494):1151–5. doi:10.1126/science.290.5494.1151.

53. Flagel LE, Wendel JF. Gene duplication and evolutionary novelty in plants. New Phytologist. 2009;183(3):557–64.

54. Flagel LE, Willis JH, Vision TJ. The standing pool of genomic structural variation in a natural population of Mimulus guttatus. Genome biology and evolution. 2013;6(1):53–64.

55. Kelly JK, Koseva B, Mojica JP. The genomic signal of partial sweeps in Mimulus guttatus. Genome biology and evolution. 2013;5(8):1457–69.

56. Mojica JP, Lee YW, Willis JH, Kelly JK. Spatially and temporally varying selection on intrapopulation quantitative trait loci for a life history trade-off in Mimulus guttatus. Molecular Ecology. 2012;21(15):3718–28.

57. Puzey JR, Willis JH, Kelly JK. Population structure and local selection yield high genomic variation in Mimulus guttatus. Molecular ecology. 2017;26(2):519–35.

58. Macnair M. A polymorphism for a lethal phenotype governed by two duplicate genes in Mimulus. Heredity. 1993;70:362–9.

59. Willis JH. Genetic analysis of inbreeding depression caused by chlorophyll-deficient lethals in Mimulus guttatus. HEREDITY-LONDON-. 1992;69:562-.

60. Bank C, Bürger R, Hermisson J. The Limits to Parapatric Speciation: Dobzhansky‒Muller Incompatibilities in a Continent-Island Model. Genetics. 2012;191(3):845–63. doi:10.1534/genetics.111.137513. PubMed PMID: PMC3389979.

61. Barton N, Bengtsson BO. The barrier to genetic exchange between hybridising populations. Heredity. 1986;57(3):357–76.

62. Gavrilets S. Hybrid Zones With Dobzhansky-Type Epistatic Selection. Evolution. 1997;51(4):1027–35. doi:10.2307/2411031.

63. Kondrashov AS, Morgan M. Accumulation of Dobzhansky-Muller incompatibilities within a spatially structured population. Evolution. 2003;57(1):151–3.

64. Lemmon AR, Kirkpatrick M. Reinforcement and the genetics of hybrid incompatibilities. Genetics. 2006;173(2):1145–55.

65. Papp B, Pal C, Hurst LD. Dosage sensitivity and the evolution of gene families in yeast. Nature. 2003;424(6945):194.

66. Doyle J, Doyle J. Genomic plant DNA preparation from fresh tissue-CTAB method. Phytochem Bull. 1987;19(11):11–5.

67. Grigoriev IV, Nordberg H, Shabalov I, Aerts A, Cantor M, Goodstein D, et al. The Genome Portal of the Department of Energy Joint Genome Institute. Nucleic Acids Research. 2012;40(D1):D26–D32. doi:10.1093/nar/gkr947.

68. Nordberg H, Cantor M, Dusheyko S, Hua S, Poliakov A, Shabalov I, et al. The genome portal of the Department of Energy Joint Genome Institute: 2014 updates. Nucleic Acids Research. 2014;42(Database issue):D26–D31. doi:10.1093/nar/gkt1069. PubMed PMID: PMC3965075.

69. Hulce D, Li X, Snyder-Leiby T, Liu CJ. GeneMarker^®^ genotyping software: tools to Increase the statistical power of DNA fragment analysis. Journal of biomolecular techniques: JBT. 2011;22(Suppl):S35.

70. Li H, Durbin R. Fast and accurate long-read alignment with Burrows‒Wheeler transform. Bioinformatics. 2010;26(5):589–95.

71. Li H, Handsaker B, Wysoker A, Fennell T, Ruan J, Homer N, et al. The sequence alignment/map format and SAMtools. Bioinformatics. 2009;25(16):2078–9.

72. Trapnell C, Roberts A, Goff L, Pertea G, Kim D, Kelley DR, et al. Differential gene and transcript expression analysis of RNA-seq experiments with TopHat and Cufflinks. Nat Protocols. 2012;7(3):562–78.

73. Kim D, Pertea G, Trapnell C, Pimentel H, Kelley R, Salzberg SL. TopHat2: accurate alignment of transcriptomes in the presence of insertions, deletions and gene fusions. Genome biology. 2013;14(4):R36.

74. Team RC. R language definition. Vienna, Austria: R foundation for statistical computing. 2000.

75. Beißbarth T, Speed TP. GOstat: find statistically overrepresented Gene Ontologies within a group of genes. Bioinformatics. 2004;20(9):1464–5.

76. Boyle EI, Weng S, Gollub J, Jin H, Botstein D, Cherry JM, et al. GO:: TermFinder—open source software for accessing Gene Ontology information and finding significantly enriched Gene Ontology terms associated with a list of genes. Bioinformatics. 2004;20(18):3710–5.

77. Guindon S, Dufayard J-F, Lefort V, Anisimova M, Hordijk W, Gascuel O. New algorithms and methods to estimate maximum-likelihood phylogenies: assessing the performance of PhyML 3.0. Systematic biology. 2010;59(3):307–21.

